# DISC1-PML protein interaction for congenital CMV infection-induced cortical neural progenitor deficit: perturbance of host signaling via viral IE1

**DOI:** 10.1101/2025.09.03.674025

**Authors:** Atsushi Saito, Stephanie Tankou, Kazuhiro Ishii, Makiko Sakao-Suzuki, Edwin C. Oh, Hannah Murdoch, Ho Namkung, Sunday Adelakun, Keiko Furukori, Masahiro Fujimuro, Paolo Salomoni, Gerd G. Maul, Gary S. Hayward, Qiyi Tang, Robert H. Yolken, Miles D. Houslay, Nicholas Katsanis, Isao Kosugi, Kun Yang, Atsushi Kamiya, Koko Ishizuka, Akira Sawa

## Abstract

Congenital CMV infection is the most common perinatal infection, affecting up to 0.5% of infants. This elicits long-term disabilities that include neuropsychiatric manifestations, such as intellectual disability, microcephaly. Despite its high prevalence, the underlying mechanism of how congenitally acquired CMV infection causes brain pathology remains unknown. Here we discovered the molecular interplay of key host (DISC1 and PML) and viral (IE1) proteins within the neural progenitor cells, which underlay an attenuated neural progenitor proliferation.

Abolishing the viral IE1 protein by delivering IE1-targeting CRISPR/Cas9 to fetal brain rescued this progenitor cell deficit, a key pathology in congenital CMV infection. A selective targeting to a viral-specific protein by the CRISPR/Cas9 system is minimal in off-target effects. Therefore, we believe that a pivotal role of IE1 in an attenuated neural progenitor proliferation in the developing cortex through its interfering with interaction between host DISC1 and PML proteins.

## INTRODUCTION

Congenital CMV infection is the most common perinatal infection, affecting up to 0.5% of infants (https://www.cdc.gov/cytomegalovirus/about/). About 10% of infected babies are symptomatic at birth, and many will suffer from permanent sequelae, including: intellectual disability, microcephaly, neuromotor deficits, and hearing loss, which remains a major medical and public health issue (1–6). The epidemiological patterns of congenital CMV infection differ according to socio-economic status (7), and CMV infection indeed exerts a pronounced burden in developing countries. For example, pregnant females suffering from infectious diseases, such as HIV, are at high risk (8–12). In the United States, the incidence of long-term sequelae from congenital CMV infection is greater than those from other perinatal complications, such as fetal alcohol syndrome and neural tube defects (13). Despite its high prevalence, the underlying mechanisms of how congenitally acquired CMV infection causes brain pathology remain unknown, resulting in limited means for prevention and treatments (1).

There have been major efforts to develop a vaccine to prevent the transmission of CMV from pregnant mothers to their offspring. Immunoglobulin therapy can decrease the severity of disabilities caused by fetal CMV infection after a primary maternal infection during pregnancy (14). However, congenital CMV infection following non-primary maternal infection accounts for a majority of congenital CMV infections (15, 16). For babies with signs of congenital CMV infection at birth, antiviral medications (e.g., ganciclovir and valganciclovir) may improve hearing and developmental outcomes. Nonetheless, a major remaining question is whether this postnatal treatment may correct the brain anomalies that happen before birth, which result in intellectual disability and long-term neuropsychiatric problems. Potential side effects of these therapies also limit their widespread application. To overcome these dilemmas, an establishment of a vaccine against CMV is awaited: a current field consensus is that a safe and effective human CMV (HCMV) vaccine is within a reach and that even a partially effective vaccine would have a major effect on the global health consequences of HCMV infection (17).

In many diseases, building animal models is an essential process to understand their pathological mechanisms, in particular when we wish their causality (18). Because of the abundance in experimental tools, mouse models are widely used (19). Indeed, to study congenital CMV infection, there are multiple mouse models (20–27). To overcome the placental barrier is refractory to CMV transmission in mice, severe combined immunodeficient (SCID) mice have been used (20). There have multiple reports that use newborn mice and built a perinatal CMV infection model (21–23). More recently, pioneered effort of delivering MCMV *in utero* has also been published, in which direct injection of MCMV into the placenta or even more directly into the fetal brains (ventricles) was made (24, 25, 27). An alternate strategy is to use guinea pig for building the model because guinea pig CMV can cross the placenta and cause infection *in utero* (28). Through these efforts, the link between CMV infection and histoanatomical/behavioral changes has been productively addressed. Nevertheless, the intracellular signaling mechanisms affected by viral infection are understudied.

In the present study, we studied an alteration in molecular signaling in neural progenitor cells (NPCs) in congenital MCMV infection. Although the prenatal influence of CMV on NPCs and its impact on a long-term disability with intellectual disability has been suggested (29–32), the molecular mechanism of the host-viral interaction remains unknown. In our present efforts, we pinned down that viral-encoded Immediate Early 1 (IE1) interferes with host protein signaling essential for proper maintenance of NPCs in the prenatal brain through the application of the CRISPR/Cas9 system against the CMV-encoded IE1.

## RESULTS

### Mouse models for congenital CMV infection

C57BL/6 mice are well characterized in many behavioral studies (33), whereas this strain is known to have CMV-resistant haplotypes associated with the natural killer cell activation in the uterus (34–36). The pioneered studies that used mouse CMV for the injection to the placenta or the fetal brains were conducted with an outbread strain ICR (Institute of Cancer Research) (25). Nevertheless, given the high utility of C57BL/6 mice because of the access to many genetically-engineered lines and the abundance of referential behavioral/neurobiological data and protocols, the use of C57BL/6 mice may be scientifically meaningful (27). Accordingly, by optimizing the timing of viral injection to minimize the resistance, we injected CMV in C57BL/6 mice at embryonic day 14.5 (E14.5), with safe delivery of pups after the living surgery.

We first used an *intraplacental injection model* (P-inj model) in which MCMV was intraplacentally injected at E14.5. Viral-encoded IE1 protein has pivotal roles in acute CMV infection by initiation of the viral replication (37, 38), and IE1 immunostaining is widely used for the evidence of CMV infection in humans and animal models (39). IE1 immuno-positive cells were observed throughout the fetal brain, when we segmented the brain in several gross regions (**Figure 1A**). In the developing cerebral cortex, most of the MCMV-infected cells were in the ventricular zone/subventricular zone (VZ/SVZ), a proliferative compartment in the developing brain (**Figure 1B**), as is evident in the human pathology (29). About 90% of IE1-positive cells inside the MCMV-infected foci in the VZ/SVZ were co-stained with Sox2, a representative marker for NPCs (**Figure S1**). Together, by adjusting the injection date to E14.5, P-inj model seems to satisfy construct and face validity at the reasonable level, with stable production and survival.

**Figure 1.**
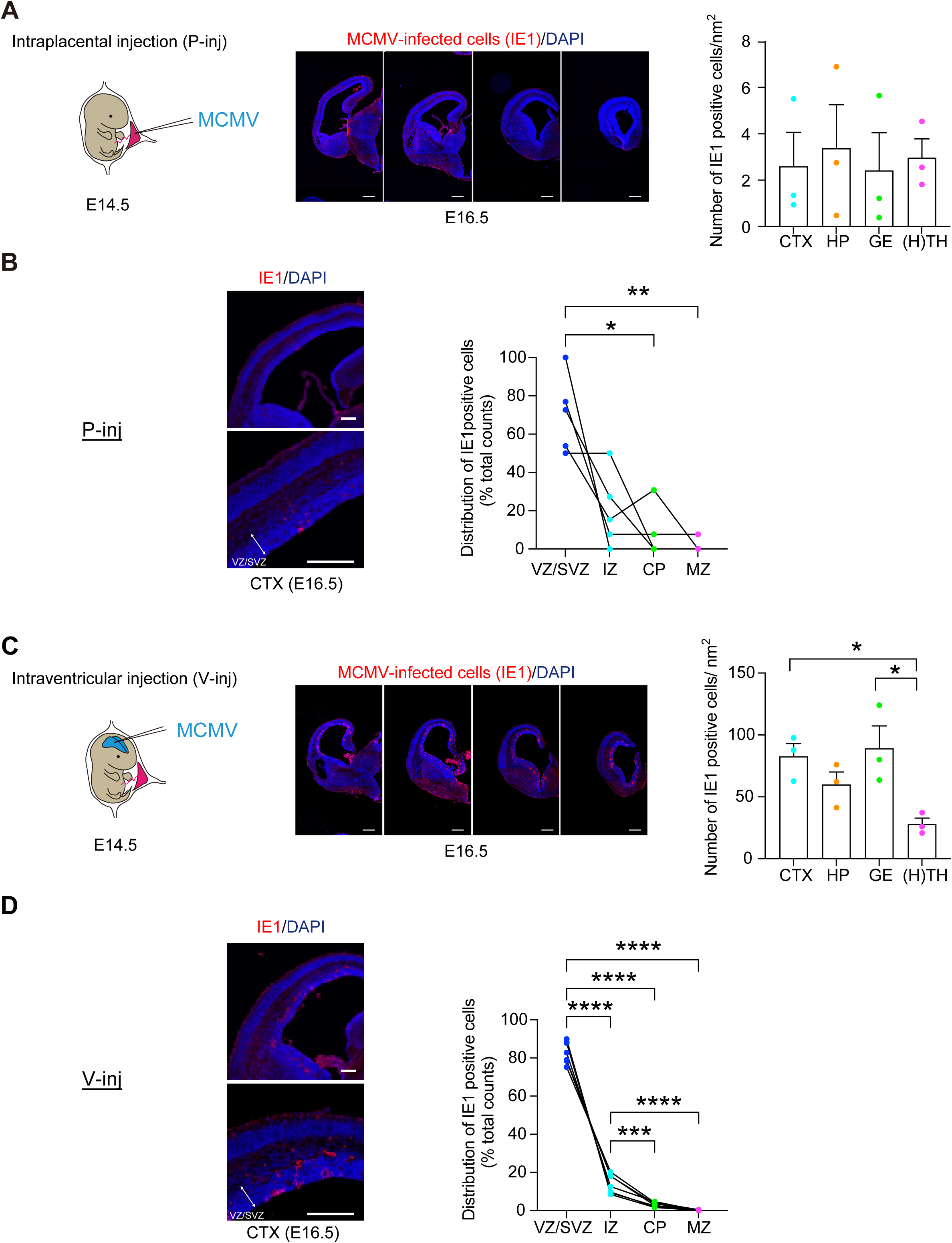
Histological validation of intraplacental (P-inj) and intraventricular (V-inj) injection models. **(A)** E14.5 embryos were intraplacentally (P-inj) injected with MCMV. Immunostaining for IE1 (red), 2 days after injection (E16.5), shows a focus of MCMV-infected cells throughout the developing brains. Sequential infected-brain images show distribution of IE1 positive cells (red) in the intraplacental (P-inj) injection models. Graphs show density of IE1 positive cells in four brain regions: cortex (CTX), hippocampus (HP), ganglionic eminence (GE), as well as thalamus and hypothalamus [H(TH)]. Scale bars, 500 µm. Graphs show mean +/-s.e.m. (n=3 mice, 12-15 slices per mice; one-way ANOVA) **(B)** Cortical distribution of IE1 cells in P-inj model. Immunostaining for IE1 (red), 2 days after injection (E16.5), shows a focus of MCMV-infected cells in the cortex. Graphs show the pattern of distribution (ratio) in the cortex: VZ/SVZ (ventricular zone/ subventricular zone), IZ (intermediate zone), CP (cortical plate), and MZ (marginal zone). In the developing cerebral cortex, most of the MCMV-infected cells were in the VZ/SVZ. The dots connected with a line indicate the data from the identical brain. Scale bars, 200 µm. (n=5 mice, \**p*<0.05, \*\**p*<0.01; one-way ANOVA) **(C)** E14.5 embryos were intracranially (V-inj) injected with MCMV. Immunostaining for IE1 (red), 2 days after injection (E16.5), shows a focus of MCMV-infected cells throughout the developing brains. Sequential infected-brain images show distribution of IE1 positive cells (red) in the intracranially (V-inj) injection models. Graphs show density of IE1 positive cells in four brain regions, indicating that IE1 positive cells appear much more densely in V-inj model compared with those in P-inj model. Furthermore, they appear more densely in CTX and GE, compared with H(TH). Scale bars, 500 µm. Graphs show mean +/-s.e.m. (n=3 mice, 12-15 slices per mice, \**p*<0.05; one-way ANOVA) **(D)** Cortical distribution of IE1 cells in V-inj model. Immunostaining for IE1 (red), 2 days after injection (E16.5), shows a focus of MCMV-infected cells in the cortex. Graphs show the pattern of distribution (ratio) in the cortex: VZ/SVZ, IZ, CP, and MZ. In the developing cerebral cortex, most of the MCMV-infected cells were in the VZ/SVZ. The dots connected with a line indicate the data from the identical brain. Scale bars, 200 µm. (n=7 mice, \*\*\**p*<0.001, \*\*\*\**p*<0.0001; one-way ANOVA)

P-inj model includes some potential limitation: the infected cells in the brain were relatively sparse, which may not be optimal when intensive characterization and intervention are performed at the cellular levels. Thus, we introduced a complementary model, in which MCMV was directly injected into the lateral ventricle of embryos at E14.5 (**Figure 1C**). Many groups including ours have utilized *in utero* gene transfer to provide plasmids to modulate gene expression in the developing cortex (40), and we applied this methodology for viral transfer. In the *ventricular injection model* (V-inj model), infected IE1-positive cells were observed throughout the fetal brain, including the cerebral cortex, which is much denser compared to those in P-inj model (**Figure 1C**). Nevertheless, these two different models (P-inj and V-inj models) showed a similar infection pattern in which most of the MCMV-infected cells were in the VZ/SVZ, a proliferative compartment in the developing brain (**Figure 1D**). Taken together, we conclude that V-inj model may be a useful alternate model to P-inj model, in particular when higher density of MCMV-infected cells is more advantageous for assays.

### NPC proliferation deficits in the developing cortex and behavioral changes in congenital CMV mouse models

The distribution of CMV in the VZ/SVZ prompted us to ask whether the viral infection might elicit defects in NPC proliferation (41), which can underlie brain anomalies and intellectual disability (42). To evaluate this possibility, by using both models, we pulse-labeled pregnant dams with bromodeoxyuridine/ 5-bromo-2’-deoxyuridine (BrdU) 2 h before euthanizing the animals on E16.5. Analysis of cortical sections revealed that BrdU signal intensity, a marker for proliferating cells, in IE1-positive cells were significantly lower than that in IE1-negative cells inside the cerebral MCMV-infected foci in both models (**Figure 2A** and **Figure S2A**). These results indicate that cellular proliferation is suppressed in IE1-positive (MCMV-infected) NPCs.

**Figure 2.**
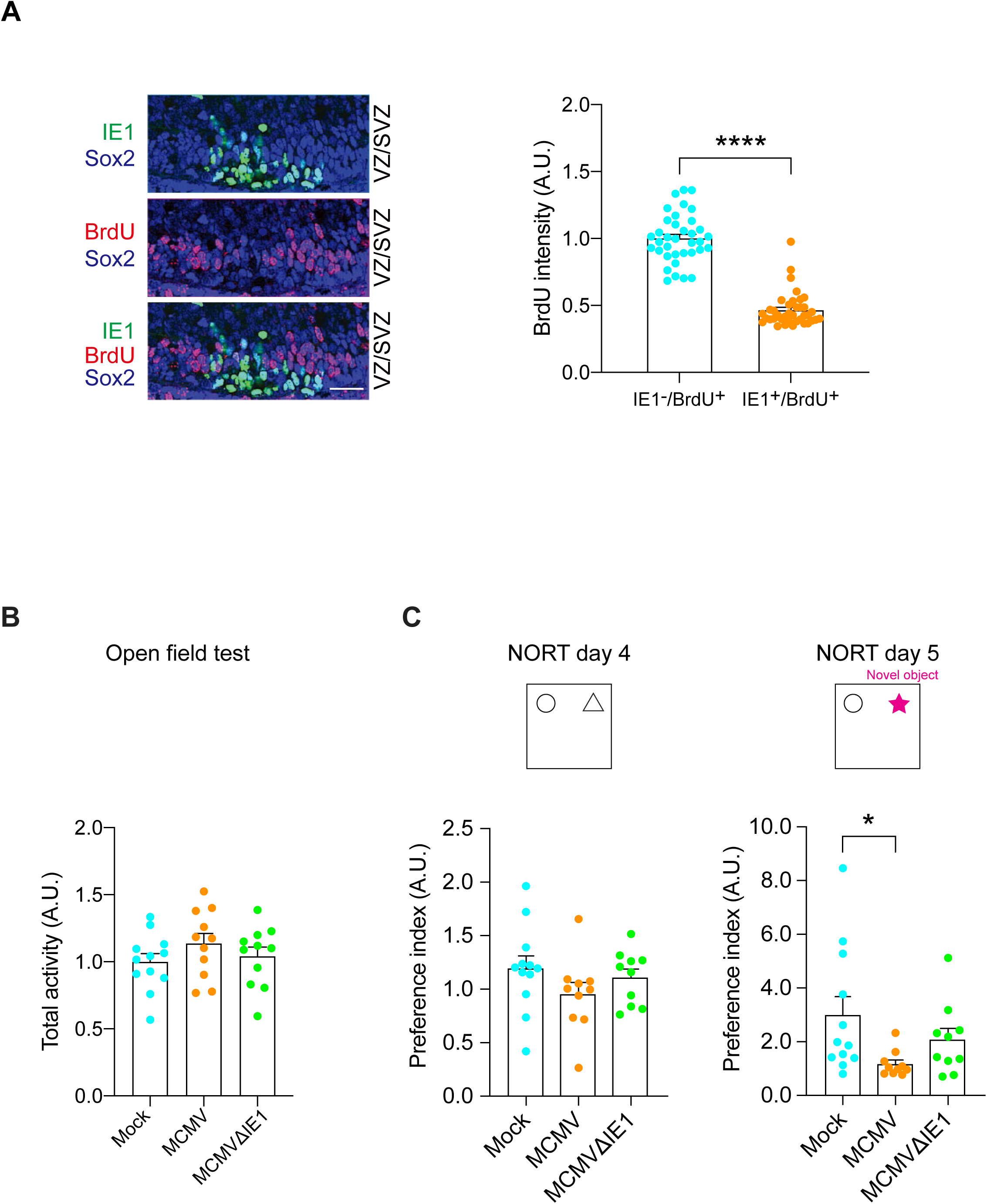
Intraplacental injection (P-inj) model shows impairment of neuronal development and a cognitive deficit. **(A)** Immunostaining for IE1 (green) and BrdU (red) shows a decrease in BrdU-labeled cells in MCMV-infected areas. IE1-positive nuclei rarely merged with BrdU-positive nuclei in P-inj model. Scale bar, 50 µm. Graph shows BrdU incorporation in uninfected NPC (Sox2-positive but IE1-negative) and infected NPC (Sox2-and IE1-double positive). The ratio of BrdU-and Sox2-double positive nuclei to total Sox2-positive nuclei, was significantly decreased in the infected area compared to the uninfected area. NPC, neural progenitor cell; Blue, Sox2; green, IE1; red, BrdU. Graph shows mean +/-s.e.m. (n=36-55 cells per group, \*\*\*\**p*<0.0001; two-sided Student’s *t* test). **(B)** The total activity of mice intraplacentally infected with Mock, MCMV, and MCMV IE1-deletion mutant (MCMV-ΔIE1) (P-inj) in the open field test. Total activity was not significantly different between groups. (Mock: n=12 MCMV: n=11, MCMVΔ1IE1: n=11; one-way ANOVA). **(C)** The results of the novel object recognition test (NORT) in three groups (Mock, MCMV, and MCMVΔIE1). Preference index (novel object exploration time / exploration time of both objects) was significantly decreased (impairment of object recognition memory) in MCMV-infected mice compared to control mice (Mock) (day5). When mice were infected with MCMVΔIE1, such impairment of object recognition memory was not observed (day 5). Graphs show mean +/-s.e.m. (Mock: n=12 MCMV: n=10, MCMVΔIE1: n=10, \**p*<0.05; one-way ANOVA).

Given that patients with congenital CMV infection often display various intellectual deficits depending on the severity of cortical lesions (43), we looked for non-specific cognitive tests relevant to various cortical dysfunction. The rodent studies using the lesion of various cortical areas have shown impaired object recognition memory measured by the novel object recognition test (NORT) (44–47). Therefore, we employed NORT to test whether a wide range of brain pathology, including the cortical pathology, was elicited by CMV infection during embryonic development. In P-inj model at postnatal day 84 (3 months of age, 3M), without differing locomotor activity, significant deficits in the NORT were observed when compared with control mice (**Figure 2B, C**). Consistent observations were also observed in V-inj model at the behavioral levels (**Figure S2B, C**). Intriguingly, mice infected with mutant MCMV that lacked IE1 (MCMVΔIE1) did not show cognitive deficits, compared with controls at the significant level (**Figure 2C**).

### CMV-encoded IE1 protein perturbs NPC proliferation by interfering with host promyelocytic leukemia (PML)-DISC1 interaction

During evolution, many pathogens develop strategies that not only enable them to survive under host conditions, but also allow them to perturb the host machinery (48). For example, human papilloma virus E6 and E7 proteins interact with host p53 and RB proteins, which leads to the disturbance of the host cell signaling cascade resulting in cervical cancer (49–52). Our experimental data that mice infected with MCMVΔIE1 exhibited no cognitive impairment may suggest that MCMV-encoded IE1 protein may interfere with a host signaling cascade, which may result in CMV infection-elicited cellular pathology in the NPCs and behavioral deficits shown above (**Figure 1** and **Figure 2**). To address this hypothesis, we ectopically overexpressed MCMV IE1 in the VZ by *in utero* gene transfer and assessed NPC proliferation. We observed a marked reduction in NPC proliferation, as indicated by BrdU labeling in the VZ/SVZ (**Figure 3A**). However, introduction of mutant IE1 (IE1Δ135-141) that lacks the domain required for disruption of PML nuclear bodies (53, 54), did not affect NPC proliferation (**Figure 3A**). These results indicate the pivotal role of MCMV-encoded IE1 in the NPC proliferation in the developing cortex, supporting the idea that viral IE1 and host PML protein interaction may be important for the pathological process.

**Figure 3.**
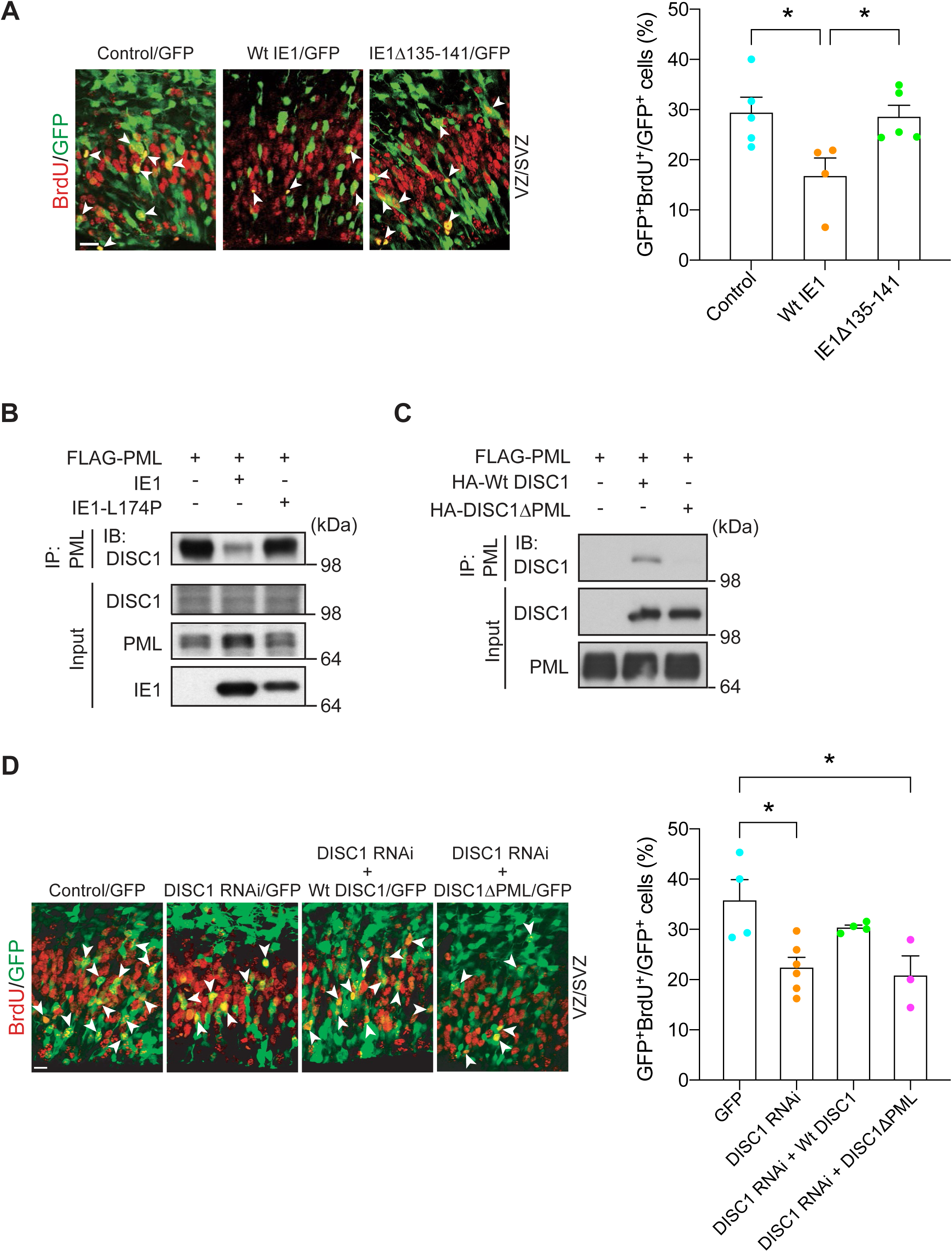
DISC1-PML interaction is required for NPC proliferation in the developing cortex. **(A)** Mouse embryos electroporated with control, wild type or mutant IE1 plasmid, and a GFP construct at E13.5 were pulse labeled with BrdU at E15.5. Green, cells transfected with GFP and control or IE1; red, BrdU-positive cells; arrowheads, GFP-and BrdU-double positive cells. VZ, ventricular zone; SVZ, subventricular zone. Scale bar, 20 µm. Bar graph represents the percentage of GFP-and BrdU-double positive cells over total GFP-positive cells in the VZ/SVZ. Graph shows mean +/-s.e.m. (Control: n=5, WtIE1: n=4, IE1Δ135-141: n=11, \**p*<0.05; one-way ANOVA). **(B)** Lysates from SK-N-MC human neuroblastoma cells co-transfected with FLAG-tagged PML and wild type IE1 or IE1-L174P were immunoprecipitated with an anti-FLAG antibody and immunoblotted with an anti-DISC1 antibody. **(C) C**os7 cells co-transfected with FLAG-tagged PML and HA-tagged wild-type DISC1, or mutant DISC1 lacking the PML binding site (DISC1ΔPML), were immunoprecipitated with the FLAG antibody and immunoblotted with the HA antibody. **(D)** Mouse embryos electroporated with various constructs at E13.5 were pulse labeled with BrdU (50 mg/kg) for 2 h at E15.5. Bar graph represents the percentage of GFP-and BrdU-double positive cells over total GFP-positive cells in the VZ/SVZ. Green, cells transfected with GFP, DISC1 RNAi, and DISC1 constructs; red, BrdU-positive cells; arrowheads indicate GFP-and BrdU-double positive cells. Scale bar, 20 µm. Graph shows mean +/-s.e.m. (GFP: n=4, DISC1 RNAi: n=6, DISC1 RNAi +Wt DISC1: n=4, DISC1 RNAi + DISC1ΔPML: n=3, \**p*<0.05; one-way ANOVA).

PML is known to function as a front-line defense against viral infection in the nuclear body (55, 56) and, reportedly, interacts with IE1 (57–59). Furthermore, in the neurodevelopmental context, DISC1 protein critically implicated in brain development and NPC proliferation is known to co-localize with the PML nuclear body (60, 61). Thus, we tested a hypothesis to see whether IE1 interferes with the DISC1-PML protein interaction. The hypothesis was experimentally validated: ectopic expression of wild-type HCMV IE1 almost completely ablates DISC1-PML binding in human cells, whereas mutant HCMV IE1-L174P, which is reportedly defective in binding with PML (57–59), failed to block the DISC1-PML protein binding (**Figure 3B**).

Consistent with these observations, infection of HCMV perturbed DISC1-PML co-localization in human cells (**Figure S3A**).

To prove that the interference of DISC1-PML binding by viral IE1 is critically relevant to CMV-elicited NPC pathology, the significance of the protein binding in proper NPC proliferation *in vivo* was investigated. We first looked for amino acid residues that are required for the binding of DISC1 with PML using established peptide array and co-immunoprecipitation approaches (62). This investigation resulted in identifying the residues 147-150 of DISC1 (**Figure 3C** and **Figure S3B**) and generating the mutant deficient of these four amino acid residues (DISC1ΔPML). As shown in previous publications (63, 64), *in utero* knockdown of DISC1 leads to NPC proliferation deficits in the developing cortex, which are rescued with co-expression of wild-type DISC1. However, we failed to observe the rescue by co-expression of DISC1ΔPML (**Figure 3D**). This suggests that DISC1-PML binding, which is disrupted by CMV-encoded IE1, is required for proper NPC proliferation during development.

### Disturbance of Notch pathway in CMV-infected NPC: a downstream of perturbed PML-DISC1 interaction

DISC1 is known to regulate NPC proliferation, at least through the Wnt pathway (63, 64). Thus, we first tested whether loss of the DISC1-PML protein interaction might affect the Wnt pathway by using β-catenin transcriptional activity as the readout in an established reporter system (63). As published by multiple groups (63–65), wild-type DISC1 fully rescued the deficits in the Wnt pathway elicited by DISC1 knockdown (**Figure 4A**). However, the effect of DISC1ΔPML was unique: although DISC1ΔPML was completely impotent of rescuing NPC proliferation defects elicited by DISC1 knockdown (**Figure 3D**), DISC1ΔPML could partially, but not fully rescue the deficits in the Wnt pathway (**Figure 4A**). One possible scenario to account for these data is that DISC1 is involved in NPC proliferation through not only the Wnt pathway but also another cascade, for both of which DISC1-PML interaction is required. *In vitro* infection study indicates that CMV dysregulates Notch pathway which may alter differentiation and proliferation in NPC (66). Thus, we hypothesized the possible involvement of the Notch pathway as the primary target. We tested this idea by using an established C-promoter binding factor 1 (CBF1) reporter system (67), because this is another critical pathway for the NPC proliferation (68). Knockdown of DISC1 suppressed Notch/CBF1 activity, which was rescued by wild-type DISC1, but not by the DISC1ΔPML mutant (**Figure 4B**). This suggests that the DISC1-PML interaction is necessary for the Notch signaling in the developing cortex. Altogether, it is likely that MCMV infection and the viral-derived IE1 protein may disturb the host DISC1-PML protein interaction, which in turn affects cellular signaling including the Notch pathway. Consistent with this notion, we observed a significant reduction in the immunoreactivity of HES1, a key transcription factor activated by Notch signaling in IE1-positive (MCMV-infected) cells compared to IE1-negative cells (**Figure 4C**).

**Figure 4.**
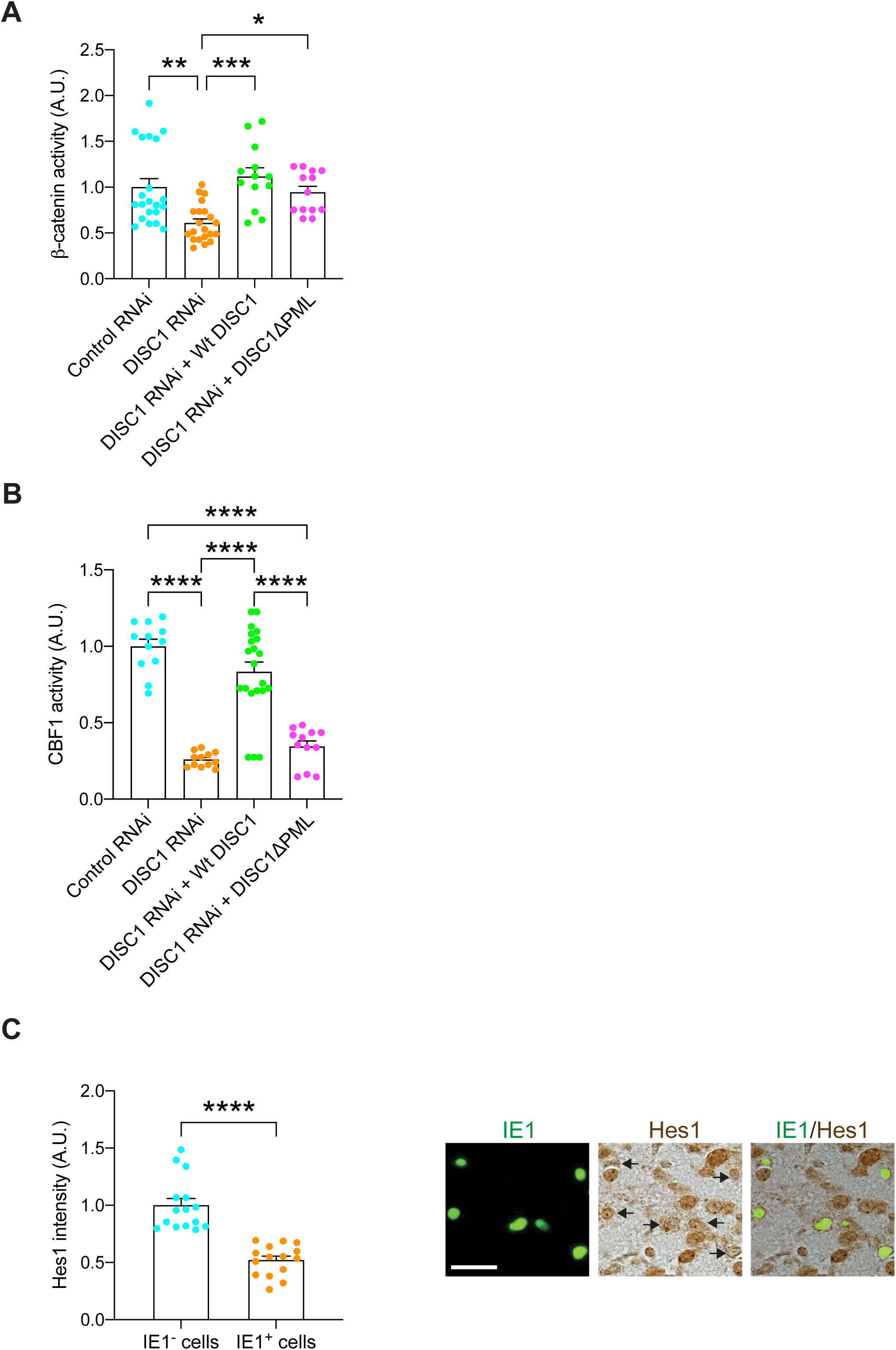
DISC1-PML protein is involved in NPC proliferation via the Notch pathway. **(A)** Super8XTOPFlash and pRL SV40 plasmids together with various constructs were injected into the lateral ventricles *in utero* at E13.5 and analyzed at E15.5. Knockdown of DISC1 suppressed β-catenin-dependent activity, which was fully rescued by wild-type DISC1 and partially rescued by DISC1ΔPML. Graph shows mean +/-s.e.m. (Control RNAi: n=21, DISC1 RNAi: n=21, DISC1 RNAi +Wt DISC1: n=13, DISC1 RNAi + DISC1ΔPML: n=13, \**p*<0.05, \*\**p*<0.01, \*\*\**p*<0.001; one-way ANOVA). **(B)** Reporter plasmids containing a CBF1-responsive element along with various constructs were injected into the lateral ventricles *in utero* at E13.5 and analyzed at E15.5. Graph shows CBF1 activity between groups and mean +/-s.e.m. (Control RNAi: n=12, DISC1 RNAi: n=12, DISC1 RNAi +Wt DISC1: n=21, DISC1 RNAi + DISC1ΔPML: n=12, \*\*\**p*<0.001; one-way ANOVA). **(C)** MCMV-infected brains were immunostained with IE1 (green) and Hes1 (brown); scale bar, 25 μm. Graph indicates the expression level of Hes1 in the IE1-positive or negative cells. Graph shows mean +/-s.e.m. (n=15 per group, \*\*\**p*<0.001; two-sided Student’s *t* test).

### CRISPR/Cas9 targeting selective to CMV-encoded IE1 to prove the significance in the NPC pathology

These data suggest that CMV-encoded IE1 is a key driver that affects host cell signaling through interfering with the DISC1-PML protein interaction, which leads to attenuated NPC proliferation, a major pathology of congenital CMV infection in the brain. Thus, targeting CMV-encoded IE1 based on the CRISPR/Cas9 system (69) and investigating the impact of the NPC pathology may provide further evidence to this notion. We introduced an all-in-one CRISPR/Cas9 vector that can express both Cas9 and the single guide RNA targeting MCMV IE1 gene. We designed target sequences by using two different web-based tools (http://chopchop.cbu.uib.no/, https://www.benchling.com/crispr/) to minimize the off-target effect, and then confirmed the successful targeting by the genome cleavage assay in MCMV infected cells (**Figure S3C**). The impact of IE1-CRISPR on the IE1 protein was confirmed by immunostaining for IE1 protein, in which the signal was significantly decreased in MCMV-infected cells co-transfected with IE1-CRISPR compared to MCMV-infected cells co-transfected with the control vector (**Figure 5A**).

**Figure 5.**
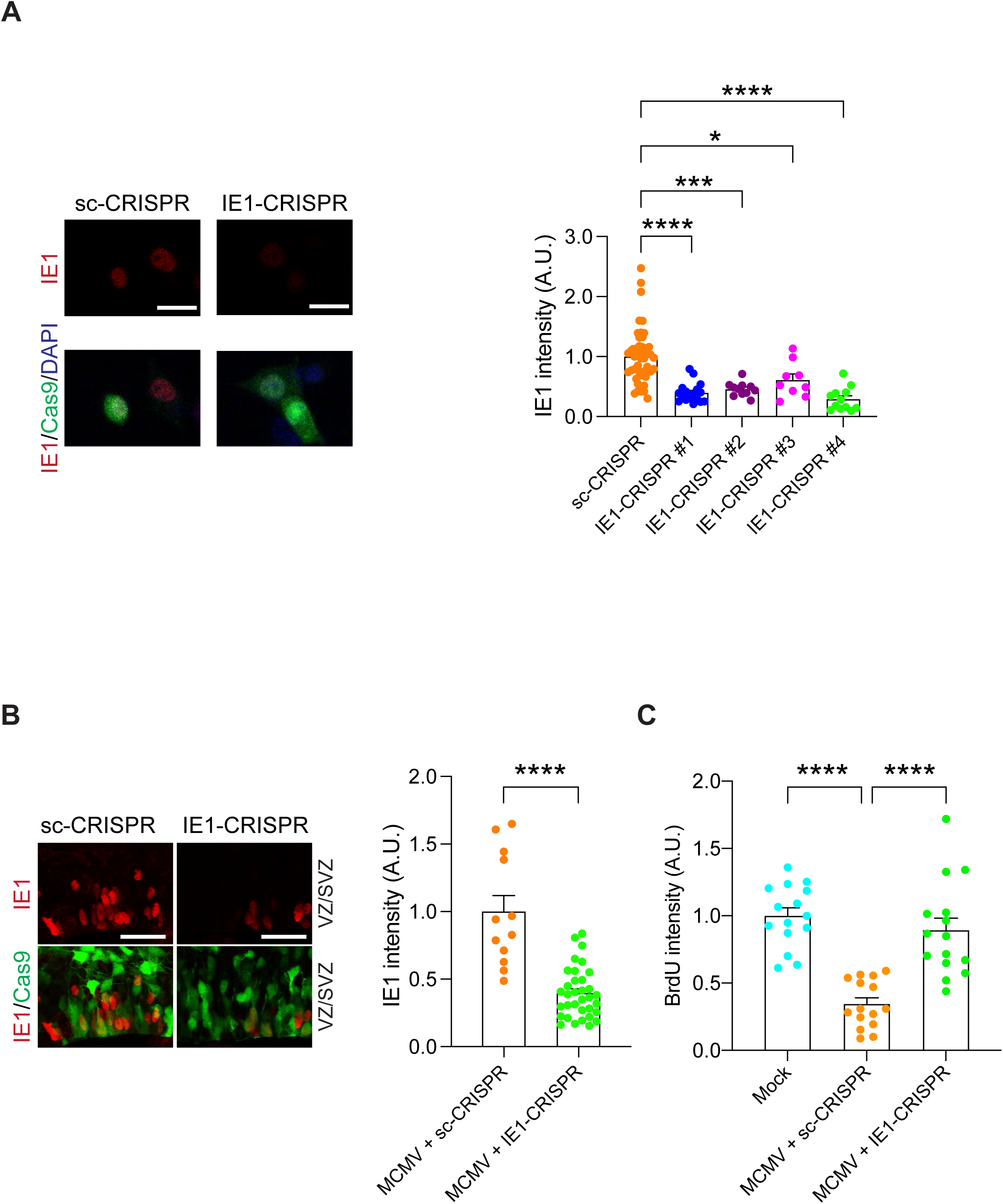
IE1-targeting CRISPR delivery via *in utero* depletes viral IE1, and rescues the brain pathologies. **(A)** HT-22 cells were transfected with IE1-CRISPR (target #1 - #4) or sc-CRISPR (control) plasmids. 24 h after transfection, MCMV was infected to the cells. Then, 24 h after infection, immunoreactivity for IE1 (red) was assessed. Graph indicates the expression level of IE1 protein in Cas9 positive cells among four candidates of IE1 target sequences and control. Red, IE1; green, Cas9; blue, DAPI. Scale bar, 25 μm. Graph shows mean +/-s.e.m. (sc-CRISPR: n=49, IE1-CRISPR#1: n=17, IE1-CRISPR#2: n=11, IE1-CRISPR#3: n=9, IE1-CRISPR#4: n=12, \**p*<0.05, \*\*\**p*<0.001, \*\*\*\**p*<0.0001; one-way ANOVA). **(B)** E14.5 embryonic brains were electroporated with IE1-CRISPR or sc-CRISPR plasmids and analyzed at E16.5. The graph indicates expression level of IE1 protein in cells transfected with IE1/or sc-CRISPR. Red, IE1; green, Cas9; scale bar, 25 μm. Graph shows mean +/-s.e.m. (MCMV + sc-CRISPR: n=12, MCMV + IE1-CRISPR: n=34, \*\*\*\**p*<0.0001; two-sided Student’s *t* test). **(C)** MCMV-infected mouse embryos electroporated with sc-or IE1-CRISPR at E14.5, and mock injected mice pulse labeled with BrdU at E16.5. The graph represents the intensity level of BrdU in IE1 and Cas9-double positive cells or cells from mock injected mice in the ventricular zone/subventricular zone (VZ/SVZ). Graph shows mean +/-s.e.m. (n=15 per group, \*\*\**p*<0.001, \*\*\*\**p*<0.0001; one-way ANOVA).

Thus, we delivered the most potent IE1-CRISPR construct (the construct #4) into the mouse fetal brain to determine its therapeutic potential for the cellular pathologies caused by congenital CMV infection. Here we used V-inj model, because the density of IE1-positive cells in P-inj model is relatively sparse, being not optimal to establish the assay condition. Two days after infection/electroporation of MCMV and the IE1-CRISPR vector into the fetal lateral ventricle, the expression level of IE1 protein was significantly decreased in brains with the IE1-CRISPR construct compared to brains with a scrambled-CRISPR (sc-CRISPR) construct (**Figure 5B**).

Consequently, the NPC proliferation deficits in MCMV-infected brains were specifically rescued by IE1-CRISPR but not by sc-CRISPR (**Figure 5C**). Taken together, our approach shows that viral IE1-targeting CRISPR delivered *in utero* diminishes MCMV-derived IE1 protein and ameliorate the NPC cellular pathology.

## DISCUSSION

In the present study, we deciphered a intracellular molecular mechanism that can account for the deficits of NPC proliferation in congenital CMV infection. We demonstrated that delivering IE1-targeting CRISPR/Cas9 to the fetal brain inhibits production of the viral IE1 protein and rescues the NPC pathology in the mouse model. The present study allows us to gain insight into a mechanism in the host pathophysiology of congenital CMV infection.

The success of modeling in C57BL/6 mice here, in contrast to many past studies that tended to use different strains to avoid the potential risk of C57BL/6-associated genetic resistance to the viral infection (25, 34, 35), will facilitate well-standardized behavioral assays and the utilization of many genetic tools available in the C57BL/6 strain for in-depth mechanistic study.

Specifically, in the present study, we uncover a mechanism for how a specific viral protein affects biological systems in the host and elicits critical pathology, which was facilitated by leveraging the delivery of the CRISPR/Cas9 system. The rapid progress of medical technology for genome editing and delivery system has provided a hope in a novel therapeutic strategy based on mechanism-guided target(s). Thus, we hope that the approach employed here may have a chance to open a new therapy of congenital CMV infection.

Given that the deficit of NPC proliferation is a key pathological event for congenital CMV infection, this question has been addressed by other groups (70, 71). For example, there is a report indicating that HCMV activates PPARγ in neural stem cells from human embryonic stem cells and brain sections from infected fetuses (70). Both PML and PPARγ are nuclear proteins, and functional cross-talks of these molecules have been reported (72). Whether and how the PML-DISC1 and PPARγ cascades are interacted will be an interesting question in the future.

We acknowledge that the present study provides many more questions that are to be addressed in future studies. First, although other rodent models may show a wider range of behavioral changes (23, 25, 73), the present models show more specific changes. This may be affected by the difference of mouse strains. We regard these differences as an opportunity of exploring the link between the neuropathology and behavioral changes by comparing different rodent models. In this case, we believe that this study providing new rodent models will be meaningful for the overall study of congenital CMV infection. Second, there is a question whether the host-viral protein interaction is only limited to NPCs or all other CMV-infected cells. Given that DISC1 is almost predominantly expressed in NPCs and early-stage differentiated neurons, this mechanism may play a role more prominently in NPCs.

## EXPERIMENTAL PROCEDURES

### Reagents and chemicals

#### Constructs

We used the RNAi constructs to mouse and human DISC1 that have been established in previous publications from other groups and ours (40, 63, 64). GST-tagged human PML IV and FLAG-tagged human PML IV deletion constructs were a generous gift from Dr.

Pandolfi (Beth Israel Deaconess Cancer Center). Human IE1 expression constructs, HA and FLAG-tagged human PML VI were obtained from Dr. Hayward (Johns Hopkins University, Baltimore, MD). Plasmid pp89 expressing MCMV IE1 expression constructs as well as mutant MCMV IE1 (IE1 with deletion of amino acids residues 135-141, IE1Δ135-141) were kindly provided by Dr. Gerd Maul (Wistar Institute, Philadelphia, PA). Target sequences in MCMV IE1 (MCMV, GenBank #U68299) were determined using the CRISPR design tool from the http://crispr.mit.edu/ (Zhang lab, MIT) and http://chopchop.cbu.uib.no (Harvard University) websites. The target oligos (**Table S1**) were inserted into the plasmid that was purchased from Addgene (pCAG-eCas9-GFP-U6-gRNA, #79145) using a restriction enzyme (BbsI) digestion and ligation approach and transformed into SURE2 competent cells (Agilent Technologies).

#### Viruses

MCMV-Smith strain and IE1(exon4)-deleted MCMV (MCMVΔIE1) and IE1(exon4)-deleted HCMV (CMVΔIE1) and the parent wild-type HCMV-Towne strain were reported previously (74, 75).

#### Antibodies

For immunofluorescence and immunoblotting, HA-tagged proteins were detected with a rat monoclonal anti-HA antibody (ROCHE) or a mouse monoclonal anti-HA antibody (Covance). Flag-tagged proteins were detected with a polyclonal antibody against FLAG (Sigma). The following antibodies were also used: mouse monoclonal anti-human PML (Santa Cruz); mouse monoclonal anti-mouse PML (Chemicon); mouse monoclonal anti-IE1 (kindly provided by Dr. Qiyi Tang, Howard University); rat anti-BrdU (Chemicon). The customized antibodies against mouse DISC1 (76) and human DISC1 (23, 61) were used to detect endogenous DISC1.

#### Recombinant proteins tagged with GST or MBP

Full-length mouse DISC1 cDNA was cloned in a pMAL vector for DISC1-MBP. The expression plasmids were introduced into Escherichia coli (E.coli) BL21, grown at 23°C with 0.1 mM IPTG. Recombinant proteins were purified from E.coli with glutathione sepharose or amylase beads.

#### Cells

Cos7 cells were grown in Dulbecco’s modified medium with 10% FBS. SK-N-MC human neuroblastoma cells were grown in DMEM/F-12 with 10% FBS. Fugene6 (Roche Applied Sciences) was used for transfection of Cos7 cells. SK-N-MC, HT-22, and HEK cells were transfected with lipofectamine 2000 (Invitrogen). SK-N-MC human neuroblastoma cells were seeded on round coverslips in 24-well plates and infected with either CMVΔIE1 or the parent Towne virus for 24 h.

### Biochemical and cellular assays

#### SPOT-synthesis of peptides and overlay experiments

Peptide libraries were produced by automatic SPOT-synthesis. Peptides were synthesized on continuous cellulose membrane supports on Whatman 50 cellulose membranes by using Fmoc (9-fluoromethyloxycarbonyl) chemistry with the AutobotSpot-Robot ASS 222 (Intavis Bioanalytical Instruments) (77).

Interaction of peptide spots with purified recombinant GST-or GST-PML (human PML IV) fusion protein was determined by overlaying the cellulose membranes with 10 mg/ml of recombinant protein. Bound recombinant proteins were detected with anti-GST antisera (Santa Cruz Biotechnologies) and a complementary HRP-coupled secondary antibody for immunoblotting.

#### Immunoprecipitation

Cells or tissues were lysed in lysis buffer (150 mM NaCl, 50 mM Tris-HCl, pH 7.5, 1% Triton X-100) containing a protease inhibitor mixture (Roche Applied Sciences). Lysates were sonicated, cell debris was cleared by centrifugation, and the soluble fraction was immunoprecipitated as described previously (40, 63).

#### Immunocytochemistry

Cells were seeded on round coverslips (Corning Glass Inc.) in 24-well plates (Falcon; Becton Dickinson Labware), washed once with phosphate-buffered saline (PBS), and fixed in ice cold methanol at-20°C for 15 min. The cells were then washed three times in PBS and blocked in 10% BSA for 30 min at room temperature. Cells were incubated with the primary antibody at 4°C overnight. After 24 h, cells were washed three times in PBS before addition of a secondary antibody conjugated with Rhodamine X (red), Cy2 or Alexa 488 (green), Cy5 (white) of either anti-rabbit, anti-rat or anti-mouse IgG for 45 min at room temperature.

After a final wash with PBS, cells were stained with DAPI.

#### In vitro genome cleavage assay

HT-22 cells were transfected with an IE1 targeting CRISPR/Cas9 plasmid or control plasmid and cultured at 37°C. 24 h after transfection, cells were infected with MCMV. 24 h after MCMV infection, cells were collected, and genomic DNA was isolated with the DNeasy^®^ Blood and Tissue Kit (QIAGEN). The region including the target sites was PCR-amplified using the primers described in **Table S2**. The PCR products were denatured and re-annealed using a thermal cycler. The heteroduplex DNA containing the insertion, deletion, or mismatched DNA (indel) was cleaved by T7 endonuclease 1. Gel analysis was performed to detect the cleaved DNA fragments.

### Animal studies - modeling and histological analyses-

All experiments were performed in accordance with the institutional guidelines for animal experiments of Johns Hopkins University and Hamamatsu University School of Medicine. The C57BL/6 male and female mice were purchased from Jackson Laboratory and used for timed-mating. These C57BL/6 mice were used for viral infection and *in utero* electroporation (see below).

#### Intraplacental MCMV injection (P-inj)

MCMV (Smith strain) was prepared and the titer of viral stock was determined as described previously. The dose of MCMV and the time of intraplacental infection and euthanization were first determined so that sufficient survival of MCMV-infected fetuses (> 80%) and adequate viral infection and BrdU incorporation in fetal brains were established. At E14.5 pregnant mice were anaesthetized by an intraperitoneal injection of 0.2–0.3 ml of 10% Somnopenthyl and uterine horns were exposed. With use of glass micropipettes, 1 x 10^5^ plaque-forming units (PFU) of MCMV in 1 µl of DMEM were injected through the uterine wall into the labyrinth region in half of the placentas. Mock-infected placentas were injected with DMEM. Two days (E16.5) after infection, BrdU (50 mg/kg) was injected intraperitoneally into pregnant mice. Three hours later, fetal brains were fixed by perfusion with 4% paraformaldehyde, immersed in 25% sucrose, and frozen in liquid nitrogen. Coronal sections were serially cut with a cryostat at 12 µm. Those days (E14.5-16.5) correspond to Carnegie Stages 20-23, late first-trimester of pregnancy in humans, are critical to brain development.

#### Ventricular MCMV injection (V-inj)

General *in utero* surgery was as described above. E14.5 pregnant mice were anesthetized, and uterine horns were exposed. With use of glass micropipettes, 1 x 10^5^ PFU of MCMV in 1 µl of DMEM were injected through the uterine wall into the lateral ventricle of embryos. Mock-infected fetuses were injected with DMEM into the lateral ventricle.

#### In utero electroporation

Pregnant C57BL/6 mice at E13.5 or E14.5 were deeply anesthetized by intraperitoneal administration of a mixed solution of Ketamine HCl (100 mg/kg) and Xylazine HCl (7.5 mg/kg), and intrauterine embryos were surgically manipulated as described previously (40, 78) with slight modifications. Plasmid solutions (1-2 μl) containing DISC1 RNAi plasmids (2 µg/µl) together with a CAG-driven GFP expression vector (1 µg/µl) were injected into the lateral ventricles. For the rescue experiments, a combination of a DISC1 RNAi plasmid (2 μg/μl in 1 μl) with a wild-type or mutant (DISC1ΔPML) DISC1 expression plasmid (1 μg/μl in 1 μl) together with a CAG-driven GFP expression vector (1 µg/µl in 1 μl) were injected. To use the same amount of DNA for each condition, an empty expression vector (pCAGGS1) was used as necessary. To perform rescue experiments in the intraplacental MCMV infection mice, an IE1 targeting CRISPR expression plasmid (1.5 μg/ul in 1 μl) together with a CAG-driven GFP expression vector (1 µg/µl in 1 μl) were injected into the lateral ventricles when the embryos were intraplacentally infected with MCMV. Electronic pulses (35 V, 50 ms, 4 times) were applied using an electroporator (CUY21E, Tokiwa Science) with a forceps-type electrode (CUY650-5, Tokiwa Science).

#### Brain slice preparation, immunohistochemistry, and BrdU incorporation assay

Histological procedures were performed as previously described with minor modifications (63). Brains were fixed with 4% paraformaldehyde, and coronal sections were obtained with a cryostat at 20 µm (CM 1850, Leica) at E16.5. For immunohistochemistry, the following primary antibodies were used: anti-BrdU (1:500) and anti-HA and anti-GFP (1:500). Fluorescent secondary antibodies conjugated to Alexa 488, Alexa 568, and Alexa 594 (Molecular Probes, Eugene, OR) were used. Nuclei were labeled with DAPI (Molecular Probes). Images of the slices were acquired with a confocal microscope (LSM 700, Zeiss). For BrdU incorporation assay, BrdU (50 mg/kg) was injected i.p. into pregnant mice 2 days after electroporation. Embryonic brains were extracted 2 h after BrdU injection. The cryosections were incubated in PBS with 0.01% Triton X100 for 30 min, and then 1N HCl at 37°C for 30 min. After washing with PBS, the sections were immunostained with a rat anti-BrdU antibody (Chemicon). Optical density of immunoreactivity in Western blotting was obtained with Image J software. Number of the IE1-positive cells was counted by the Volocity software (Parkin Elmer).

#### β-catenin and CBF1 activity assays with luciferase reporter system

Luciferase reporter system assays were carried out by using either the β-catenin or CBF1 reporter plasmid together with pRL-SV40 plasmid. Expression and/or RNAi constructs, as well as reporter plasmids, were delivered into the ventricular zone at E13.5 by *in utero* electroporation. The luciferase activity was measured at E15.5.

#### Behavioral assays

Only male mice were used for the behavioral assays to avoid confounding effects from the female estrous cycle. Open field test was conducted as described previously (79) with minor modifications. In brief, each mouse was placed in a transparent acrylic cage (40 cm x 40 cm; San Diego Instruments, San Diego, CA) for 30 min. The test was Locomotor activities were recorded by an infrared activity monitor (San Diego Instruments). A single beam break was reported as “count”. For novel object recognition test, mice were habituated to the testing box over 3 days. On day 4, the mice were exposed to two objects for 10 min. On day 5 (retention), one familiar object was replaced with a novel object and the mice were allowed to explore both objects for 5 min. The preference index was calculated as the ratio of time spent exploring the novel object in the retention session to total exploration time.

### Statistical analyses

All statistical analyses were performed using GraphPad Prism 10. For comparisons between two independent groups, two-tailed Student’s t-tests were used, with statistical significance defined as *p* < 0.05. For comparisons involving more than two groups, one-way ANOVA was conducted, followed by Tukey’s post hoc test to assess pairwise group differences. Results were considered statistically significant if *p* < 0.05 based on Tukey’s test.

## DATA AVAILABILITY

All data generated or analyzed during this study are included in this article and its supplementary information files.

## SUPPORTING INFORMATION

This article contains supporting information.

## ACKNOWLEDGMENTS AND FUNDING INFORMATION

We thank Ms. Ruth Cruz for virus preparation, Ms. Y. Lema for preparing the figures and organizing the manuscript, and Dr. M. Salgado, Dr. P. Talalay and Ms. L. Guttman for critical reading of this manuscript. This work was supported by USPHS grants MH136297 Silvo O. Conte center (A. Sawa, A. Kamiya, and K. Yang), MH094268 Silvo O. Conte center (A. Sawa, N. Katsanis, and A. Kamiya), MH105660 (A. Sawa and K. Ishizuka), MH091230 (A. Kamiya), MH96208 (K Ishizuka), SC1AI112785 (Q. Tang) as well as foundation grants from the Stanley Foundation (A. Sawa), RUSK/ S&R Foundation (A. Sawa), Brain & Behavior Research Foundation (A. Saito, E. E. Oh, K. Ishizuka, A. Kamiya and A. Sawa), Japan Health Sciences Foundation (SHC4401; I. Kosugi), and Subsidies for Current Expenditures to Private Institutions of Higher Education from the Promotion and Mutual Aid Corporation for Private Schools of Japan (K Ishizuka).

## AUTHOR CONTRIBUTIONS

A. Saito: Methodology, Investigation, Writing - Original Draft; S. Tankou: Methodology, Investigation; K. Ishii: Methodology, Investigation; M. Sakao-Suzuki: Investigation; E. C. Oh: Investigation; H. Murdoch: Investigation; H. Namkung: Investigation; S. Adelakun: Investigation; K. Furukori: Investigation; M. Fujimuro: Investigation; P. Salomoni: Resources; G. G. Maul: Resources; G. S. Hayward: Resources; Q. Tang: Resources; R. H. Yolken: Investigation; M. D. Houslay: Investigation; N. Katsanis: Investigation; I. Kosugi: Investigation; K. Yang: Formal analysis; A. Kamiya: Investigation; K. Ishizuka: Methodology, Investigation, Writing - Original Draft; A. Sawa: Methodology, Conceptualization, Supervision, Project administration, Funding acquisition, Writing - Original Draft.

