## Supplemental information for "DISC1-PML protein interaction for congenital CMV infection-induced cortical neural progenitor deficit: perturbance of host signaling via viral IE1"

**Figure S1**

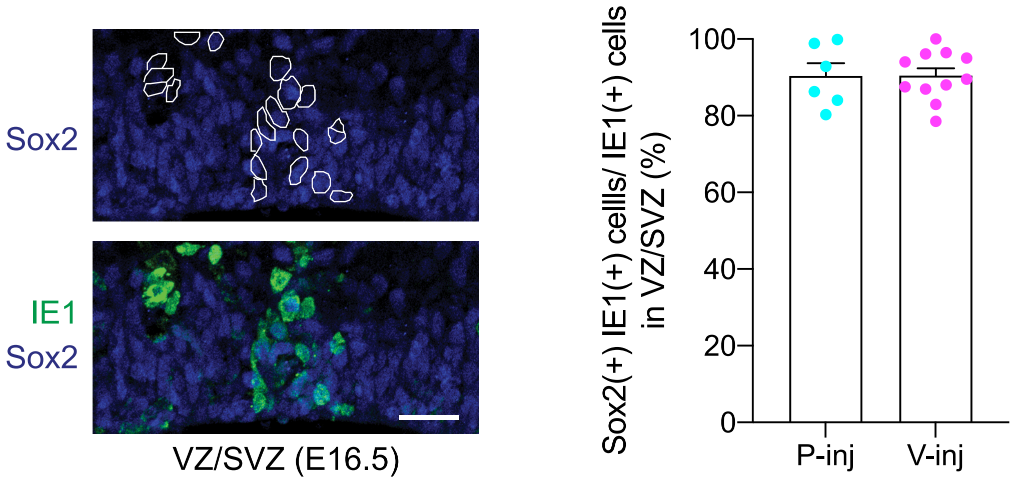
Sox2
IE1
Sox2
VZ/SVZ (E16.5)
Sox2(+) IE1(+) cells/ IE1(+) cells in VZ/SVZ (%)
100
80
60
40
20
0
P-inj
V-inj

**Figure S1.** IE1 highly localizes in Sox2 positive neural progenitor cells (NPCs) (related to Fig. 1).

Immunostaining for IE1 (green) and Sox2 (blue) in the ventricular zone/ subventricular zone (VZ/SVZ). The white lines indicate the shapes of IE1 immunoreactivity. Scale bar, 50 µm. Graph shows percentage of Sox2 positive cells in IE1 positive cells (mean +/- s.e.m, n=6-11 images). There were no differences in the two models (*P*=0.9805; two-sided Student's *t* test).

**Figure S2**

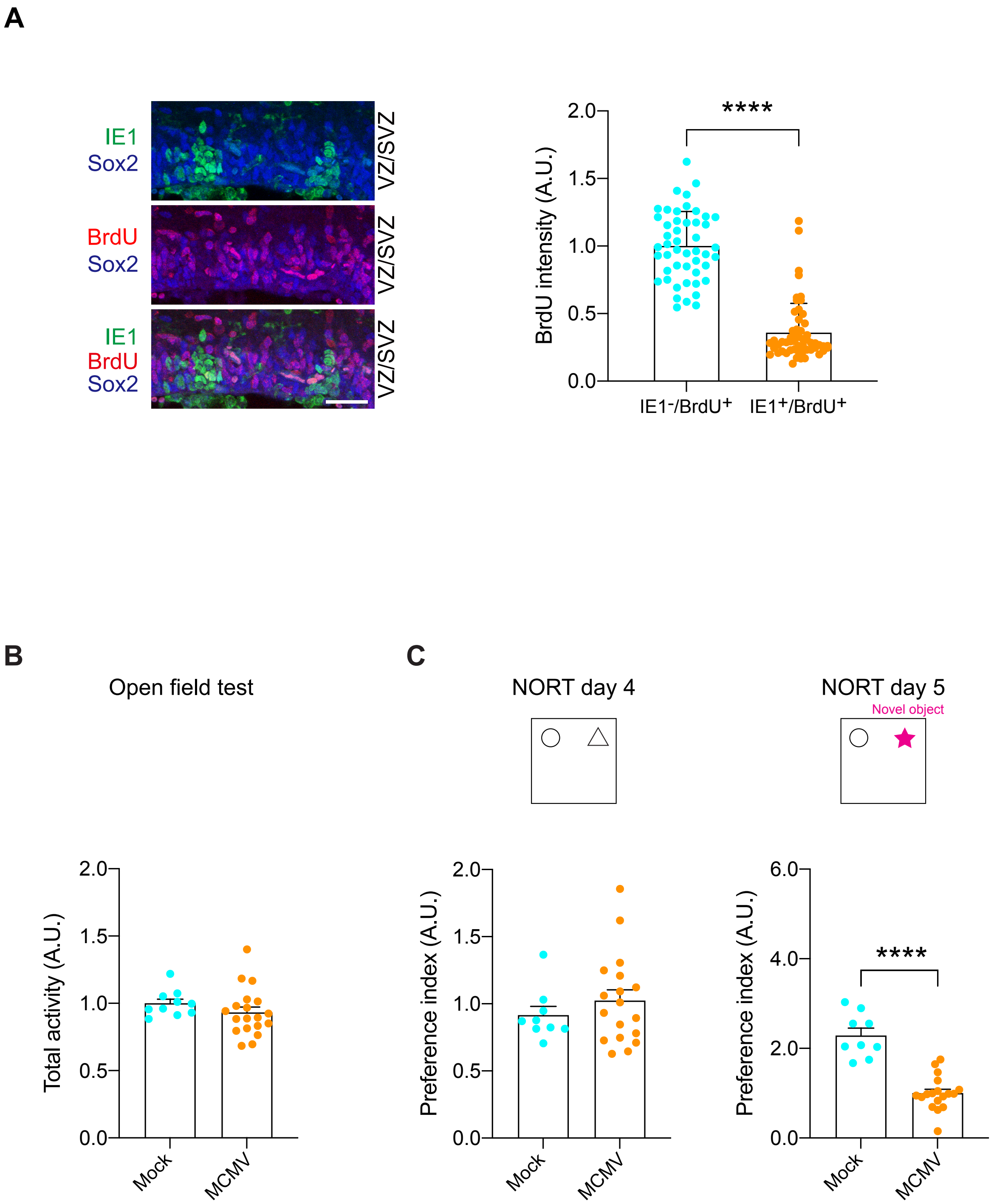
A
IE1
Sox2
BrdU
Sox2
IE1
BrdU
Sox2
VZ/SVZ
VZ/SVZ
VZ/SVZ
BrdU intensity (A.U.)
2.0
1.5
1.0
0.5
0.0
****
IE1-/BrdU+
IE1+/BrdU+
B
Open field test
Total activity (A.U.)
Mock
MCMV
C
NORT day 4
NORT day 5
Novel object
Preference index (A.U.)
6.0
4.0
2.0
0.0
****

**Figure S2.** Intraventricular injection (V-inj) model shows impairment of neuronal development and a cognitive deficit (related to Fig. 2).

**(A)** Immunostaining for IE1 (green) and BrdU (red) shows a decrease in BrdU-labeled cells in MCMV-infected areas. IE1-positive nuclei rarely merged with BrdU-positive nuclei in the V-inj model. Scale bar, 50 µm. Graph shows BrdU incorporation in uninfected NPC (Sox2-positive but IE1-negative) and infected NPC (Sox2- and IE1- double positive). The ratio of BrdU- and Sox2- double positive nuclei to total Sox2-positive nuclei was significantly decreased in the infected area compared to the uninfected area. NPC, neural progenitor cell; Blue, Sox2; green, IE1; red, BrdU. Graph shows mean +/- s.e.m. (n=36-55 cells per group, ****$P$<0.0001; two-sided Student's *t* test).

**(B)** The total activity of mice infected with Mock and MCMV in the open field test. Total activity was not significantly different between groups. (Mock: n=10, MCMV: n=19; two-sided Student's *t* test).

**(C)** The results of the NORT in two groups (Mock and MCMV) in the ventricular injection (V-inj) model at 1 month after infection. Graphs indicate the preference index among the groups on day 4 and day 5 of testing. The preference index was calculated as the ratio of exploration time for novel object (or right object on day 4)/exploration time for both objects.

Graphs show mean +/- s.e.m. (Mock: n=9, MCMV: n=18, ***$P$<0.001, ****$P$<0.0001; two-sided Student's *t* test).

### Figure S3

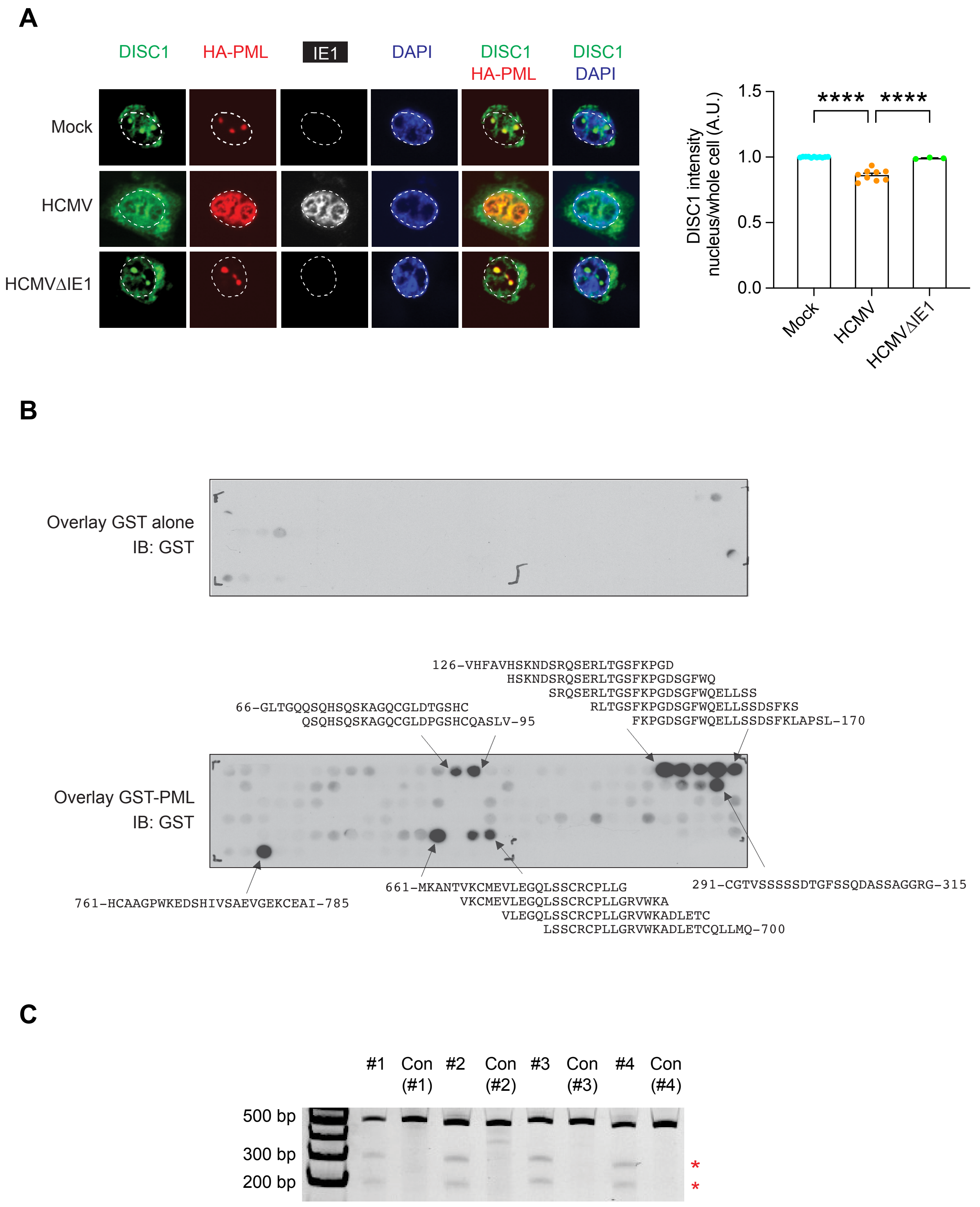
A
DISC1
HA-PML
IE1
DAPI
DISC1 HA-PML
DISC1 DAPI
Mock
HCMV
HCMVΔIE1
DISC1 intensity nucleus/whole cell (A.U.)
1.5
1.0
0.5
0.0
****
****
Mock
HCMV
HCMVΔIE1
B
Overlay GST alone IB: GST
126-VHFAVHSKNDSRQSERLTGSFKPGD
HSKNDSRQSERLTGSFKPGDSGFWQ
SRQSERLTGSFKPGDSGFWQELLSS
RLTGSFKPGDSGFWQELLSSDSFKS
FKPGDSGFWQELLSSDSFKLAPSL-170
66-GLTGQQSQHSQSKAGQCGLDTGSHC
QSQHSQSKAGQCGLDPGSHCQASLV-95
Overlay GST-PML IB: GST
761-HCAAGPWKEDSHIVSAEVGEKCEAI-785
661-MKANTVKCMEVLEGQLSSCRCPLLG
VKCMEVLEGQLSSCRCPLLGRVWKA
VLEGQLSSCRCPLLGRVWKADLETC
LSSCRCPLLGRVWKADLETCQLLMQ-700
291-CGTVSSSSSDTGFSSQDASSAGGRG-315
C
#1
Con (#1)
#2
Con (#2)
#3
Con (#3)
#4
Con (#4)
500 bp
300 bp
200 bp
*
*

**Figure S3.** *In vitro* validations (related to Fig. 3 and Fig. 5)

**(A)** Co-localization of DISC1-PML in HCMV-infected cells. Images show the nuclei of neuroblastoma cells. Infection with HCMV disturbed PML-DISC1 co-localization in human neuroblastoma. However, infection of HCMV lacking IE1 (HCMV ΔIE1) did not affect the PML-DISC1 co-localization. Green, DISC1; red, HA-PML; white, IE1; blue DAPI. Graph shows mean +/- s.e.m. (Mock: n=11, HCMV: n=9, HCMVΔIE1: n=3, *$P<0.05$, **$P<0.001$; one-way ANOVA).

**(B)** Pinpointing the PML binding domain on DISC1. DISC1 peptide arrays were probed for PML interaction sites. The dot blot shows interaction between GST-PML and a DISC1 peptide array.

**(C)** Viral IE1-targeting CRISPR/Cas9 cleaves the IE1 gene. The T7 Endonuclease I assay, which can detect genome cleavage caused by CRISPR/Cas9, shows that the IE1 genes in the MCMV genome are cleaved in MCMV-infected HT22 cells when co-transfected with the IE1-targeting CRISPR construct (IE1-CRISPR). Con, control. Red asterisks indicate cleavage products.

**Table S1.** Single guide RNA target sequences.

| Target | Sequence | PAM sequence |
|---|---|---|
| IE1-CRISPR#1 | 5'- CGGCACGCTCATCTAGTGCG -3’ | TGG |
| IE1-CRISPR#2 | 5'- AGATTAGTGGGCATGAAGTG -3’ | TGG |
| IE1-CRISPR#3 | 5'- CACTAGATGAGCGTGCCGCA -3’ | TGG |
| IE1-CRISPR#4 | 5'- GATGCGCTCGAAGATATCAT -3’ | TGG |

**Table S2.** T7 endonuclease I assay primer sequences.

| Target | Primer |
|---|---|
| IE1-CRISPR#1 | Fw: 5'- GATATCTTCGAGCGCATCGA -3'<br>Re: 5'- ACACACCCTGTGATATTGG -3' |
| IE1-CRISPR#2 | Fw: 5'- CTGTTGTCCTGTAAGATTGC -3'<br>Re: 5'- CTTCCACCACTACCACATG -3' |
| IE1-CRISPR#3 | Fw: 5'- TCGAAAGACAACGCAAGATG -3'<br>Re: 5'- GGTCTCTAGATGGTCTTTCATG -3' |
| IE1-CRISPR#4 | Fw: 5'- CTCAGTTATTCACATCATGAC -3'<br>Re: 5'- TGTAACAGGGTGGATCATG -3' |
